# Characterization of RNA content in individual phase-separated coacervate microdroplets

**DOI:** 10.1101/2021.03.08.434405

**Authors:** Damian Wollny, Benjamin Vernot, Jie Wang, Maria Hondele, Anthony Hyman, Karsten Weis, J. Gray Camp, T.-Y. Dora Tang, Barbara Treutlein

**Affiliations:** Max Planck Institute for Evolutionary Anthropology, Leipzig, Germany; Max Planck Institute of Molecular Cell Biology and Genetics, Dresden, Germany; Cluster of Excellence Physics of Life, TU Dresden, Dresden, Germany; Institute of Biochemistry, ETH Zurich, Zurich, Switzerland; Institute of Molecular and Clinical Ophthalmology, Basel, Switzerland; Department of Biosystems Science and Engineering, ETH Zürich, Basel, Switzerland; Present address: RNA Bioinformatics and High Throughput Analysis, Friedrich Schiller University Jena, Germany; Present address: Biozentrum, University of Basel, Basel, Switzerland

## Abstract

Liquid-liquid phase separation or condensation is a form of macromolecular compartmentalization. Condensates formed by complex coacervation were hypothesized to have played a crucial part during the origin-of-life. In living cells, condensation organizes biomolecules into a wide range of membraneless compartments. Although RNA is a key component of condensation in cells and the central component of the RNA world hypothesis, little is known about what determines RNA accumulation in condensates and how single condensates differ in their RNA composition. Therefore, we developed an approach to read the RNA content from single condensates using high-throughput sequencing. We find that RNAs which are enriched for specific sequence motifs efficiently accumulate in condensates. These motifs show high sequence similarity to short interspersed elements (SINEs). We observed similar results for protein-derived condensates, demonstrating applicability across different in vitro reconstituted membraneless organelles. Thus, our results provide a new inroad to explore the RNA content of phase-separated droplets at single condensate resolution.

## Introduction

In the 1920’s de Jong coined the term coacervation to describe a liquid-liquid phase separation process between two oppositely charged polymers in solution^1^. Electrostatic interactions bring the two components together, subsequent entropic release from water and counter ions from around the polyelectrolytes drive phase separation into membrane-free and chemically enriched micron-sized droplets^2^. These coacervate droplets have been shown to form from a wide variety of different molecules with very little chemical specificity from synthetic polyelectrolytes, to biological polyelectrolytes and small charged molecules^3^. Consequently, they were hypothesized to play a role in the origin-of-life by bringing together the first molecules to spatially localize the first primitive reactions^4^. Since then coacervates formed from synthetic polymers have been exploited in a range of industries from food separation to pharmaceuticals^5^. More recently, it has been shown that the coacervation process plays an active role in the liquid-liquid phase separation of condensates in biological systems. Whilst, the mechanism of formation of biomolecular condensates in cells has now been extensively studied, an understanding of how condensates regulate biochemical processes in time and space is still in its infancy^6, 7^.

Key to unravelling these unanswered questions is deconvoluting the molecular content and physicochemical properties of the condensates. So far, progress in this area has been limited by difficulty in isolating condensates from cells in their dynamic environment. To this end, *in vitro* reconstitution has been instrumental for in depth droplet characterization^8^.

Most of the condensate characterization has relied on fluorescent microscopy. Indeed, characterization of the partition coefficients has only recently been optimized using high-throughput microfluidic methods based on fluorescence of single solutes^9^. Despite this progress, there remains no methodology to determine the precise amount and type of molecules in individual coacervate droplets. To this end, we have exploited single-cell RNA sequencing technology and developed a novel way to determine the amount and sequence of RNA incorporated into individual coacervate droplets. This provides an unprecedented opportunity to determine, for the first time, the RNA content of individual coacervate droplets within a population. Furthermore, we show how this method can be applied to both synthetic coacervate microdroplets and condensates prepared from biological phase separating protein scaffolds such as the human RNA-binding protein Fused in Sarcoma (FUS) and yeast DExD/H-box helicase 1 (Dhh1). We identify the RNA properties that are crucial for uptake into synthetic coacervates and demonstrate similarity to FUS and Dhh1 droplets depending on the coacervate chemical identity. This provides for the first time a direct link between synthetic coacervates and biomolecular condensates in cells, implying that coacervates can serve as models of biological systems.

## Results

In order to determine the RNA content of individual coacervate droplets or condensates, we aimed to work with the following four droplet systems: carboxylmethyldextran (CM-Dex) and Poly(diallyldimethylammonium chloride) (PDDA) (molar ratio: 6:1) or CM-Dex with polylysine (pLys) microdroplets (molar ratio: 6:1) in 10 mM Tris and 4mM MgCl_2_ at pH 8 or recombinant FUS (25 mM Tris-HCL, 150 mM KCL, 2.5 % glycerol, 0.5 mM DTT, pH 7.4) or recombinant Dhh1 (50 mM KCl, 30 mM HEPES-KOH, 2 mM MgCl_2_, pH 7.4) condensates were prepared in the presence of total RNA (50 ng/μl). The total RNA was isolated from human induced pluripotent stem cells (iPSCs) immediately before each experiment.

We started by analyzing the RNA content of CM-Dex:PDDA coacervates (Fig. 1a). The RNA-containing membrane-free droplets were loaded into 96 well plates with each well containing 4 μL of guanidine hydrochloride (6 mM) by fluorescence activated cell sorting (FACS). Using this technique, it was possible to precisely control the number of droplets in each well - down to single coacervates. The presence of high concentration guanidine hydrochloride led to a change in turbidity of the coacervate dispersion from cloudy to clear which is synonymous with the dissolution of coacervate droplets (Supplementary Fig. 1a). This indicates that the coacervate droplets are dissolved upon addition to the well plate releasing the RNA from the droplets. Droplets sorted into well plates were immediately frozen at −80 °C. The released RNA was purified by magnetic solid phase reversible immobilization (SPRI) beads. The magnetic beads were added to a buffer containing dNTP/oligodT for reverse transcription of messenger RNA (mRNA) to generate complementary DNA (cDNA). We focused our analysis on mRNA (commonly referred to as transcripts) because of its heterogeneity in terms of sequence composition and length providing us with data from a pool of highly diverse RNAs.

**Fig. 1.**
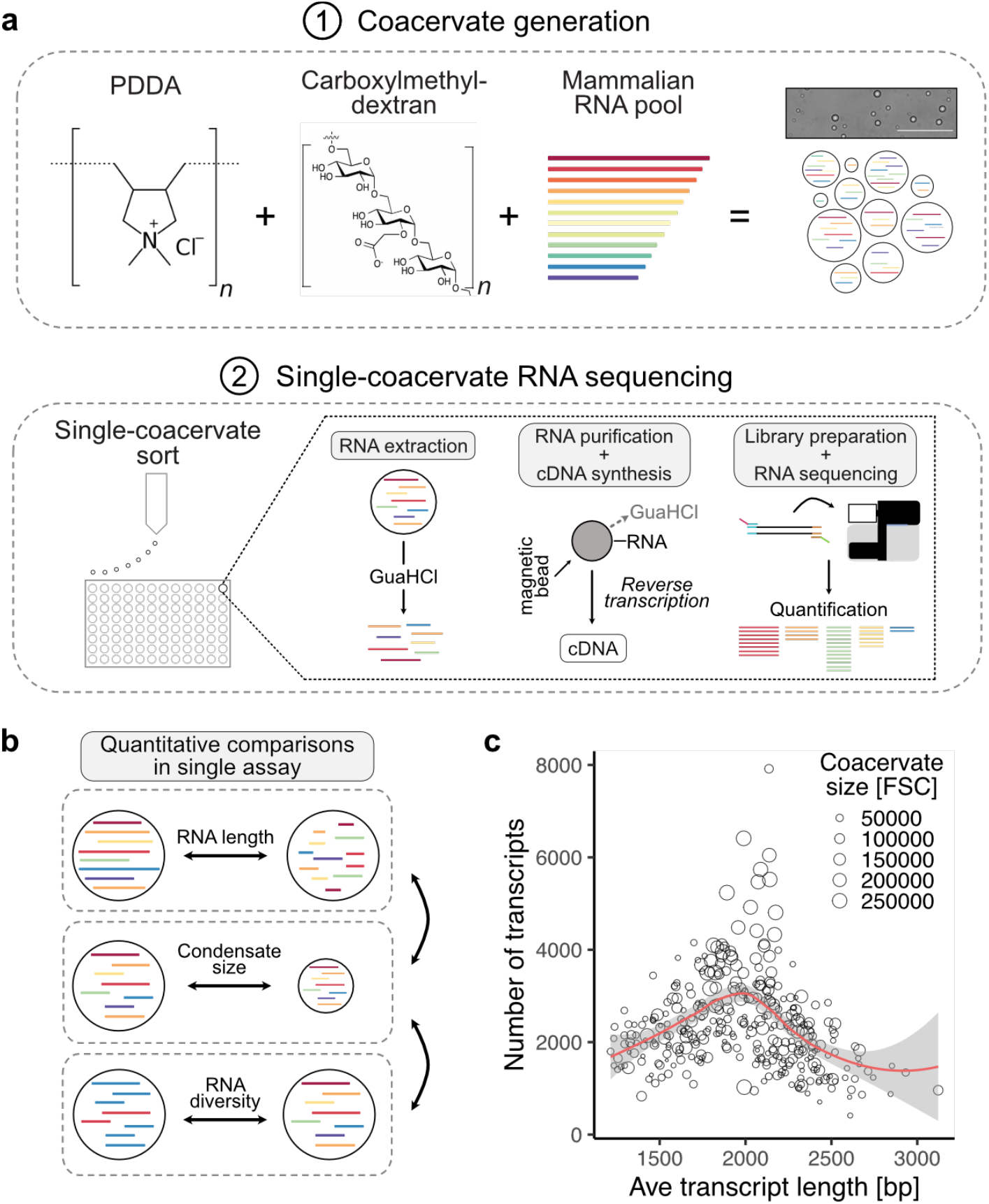
Sequencing RNA of single phase-separated coacervates. **a**, Schematic of coacervate generation and single-coacervate sequencing strategy. Coacervates were generated by mixing Carboxymethyldextran with PDDA (CM-Dex:PDDA). Total RNA isolated from iPS was used as RNA input. Single coacervates were sorted into 96 well plates using fluorescence-activated cell sorting (FACS). RNA was extracted from each coacervate and mRNA was converted to cDNA and sequenced upon library preparation. RNAs present in each sequenced coacervate were computationally identified and quantified. **b**, Schematic illustration of cross comparisons of RNA length, coacervate size and complexity of RNA pool from hundreds of individual coacervates. **c**, Relationship between the size of single coacervates, the number of different RNA transcripts and the average length of all RNA transcripts in each coacervate. Each dot represents a sequenced coacervate. Coacervate size was measured by the FACS forward-scatter (FSC).

Full length cDNA was amplified by PCR as previously described^10^ (Fig. 1a). Illumina sequencing of the RNA content of individual coacervates showed that it was possible within the resolution of the experiment to determine the sequence, length and relative abundance (transcripts per million, TPM) by computationally matching the sequenced RNA fragments to known reference RNA sequences of the human genome. In addition, the relative size of the individual droplets was obtained from FACS by the forward scatter of the droplet. Together with sequence analysis, it is possible for the first time to obtain information on both the genotype and phenotype within a population of coacervate microdroplets on a single droplet level.

Bioanalyzer traces were used for quantification and quality control of the amplified cDNA from 0, 1, 10, 100 and 1000 CM-Dex:PDDA coacervate droplets and demonstrated successful library preparation even from single coacervates (Supplementary Fig. 1b). Furthermore, quantification of the amount of amplified cDNA showed a linear correlation to the amount of RNA with increasing number of coacervate droplets (Supplementary Fig. 1c). Even though errors in quantification may arise from PCR amplification steps, these results indicate that the methodology of extraction and amplification of RNA is robust and consistent.

Using this approach, we aimed to investigate the relationship between coacervate size and its RNA content - specifically the relationship between the diversity of RNA transcripts, the average length of the transcripts within the coacervates and the coacervate size (Fig. 1b). Our results showed that the largest coacervates had the highest diversity of transcripts (Fig. 1c). In comparison coacervates containing the longest average transcript length were among the smallest coacervates. These smaller coacervates also displayed a very low diversity of detected RNA transcripts (Fig. 1c). Interestingly these results indicate that random pools of RNA will localize in a heterogeneous nature within dispersions of coacervate droplets leading to different phenotypic properties.

Next, we wanted to test if the RNA distribution within the coacervate population was consistent across experiments. We found that the frequency with which transcripts are detected in condensates is highly reproducible across experiments (Pearson correlation coefficient r = 0.86, Fig. 2a). This indicates that, albeit being a dynamic process, the localization of RNA into condensates is not random. Interestingly, the relative amount of each transcript within the condensates is not as consistent between experiments as the frequency with which specific transcripts are detected across condensate (r = 0.58, Fig. 2b). Furthermore, the correlation for the relative amount of each RNA remained low for both small and large droplets (Supplementary Fig. 2). Whilst these results show that the experiments are reproducible for the type of RNA, every coacervate dispersion produced in the presence of random RNA will lead to a different heterogeneous population with respect to the relative amount of RNA. This has very interesting implications in considering the role of coacervation in origin-of-life and modern biological studies where each droplet within a pool may have different genotypic properties.

**Fig. 2.**
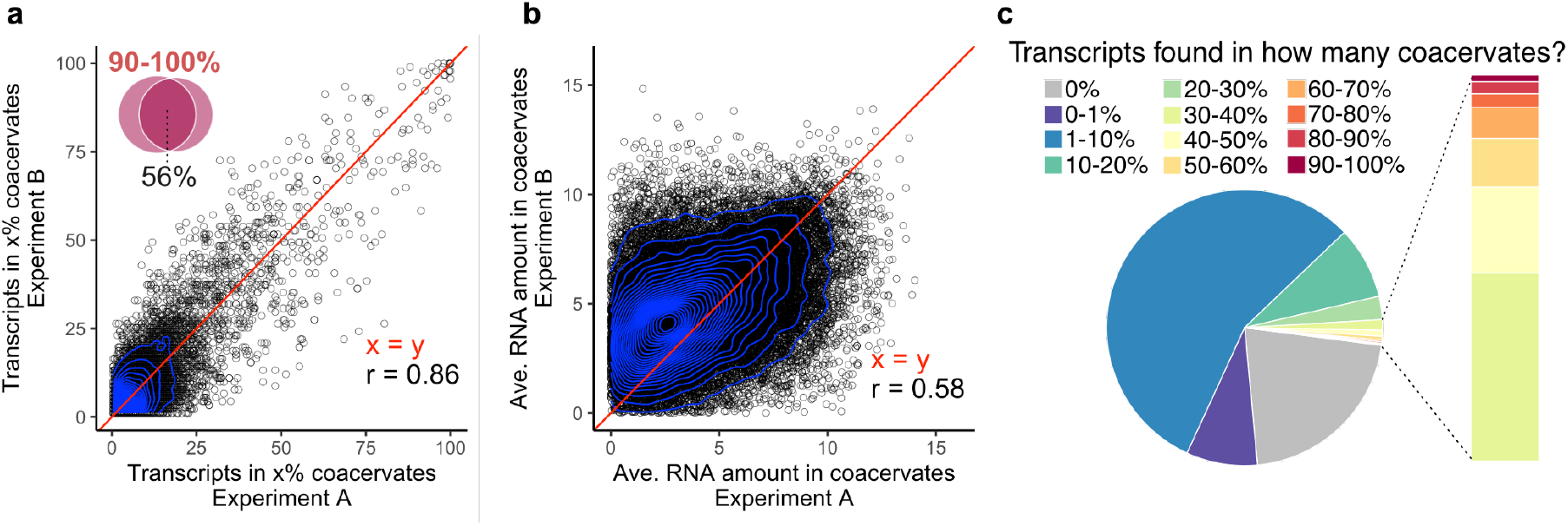
Comparison of experiment-to-experiment variability of RNA detection in coacervates. **a**, Quantification of the efficiency of RNA assembly into CM-Dex:PDDA coacervates across independent experiments. Each dot represents an RNA transcript. Venn diagram: Overlapping transcripts across experiments that were found in 90-100% of coacervates. **b**, Experiment-to-experiment variation of the average abundance of each RNA transcript across all coacervates in which it was detected. RNA abundance for each transcript is calculated as transcripts per kilobase million (log2(TPM)) enabling comparison of relative transcript abundances across coacervates. Red line indicates perfect correlation (x = y). Pearson correlation coefficient = r, **c**, Pie chart demonstrating how frequently each input RNA transcript was detected in coacervates.

We further quantified the relative percentage of each input RNA transcript in CM-Dex:PDDA coacervates (Fig. 2c). This would allow us to characterize the proportion of input transcripts that were found within the dispersion of the droplets. Our analysis showed three things: 1. We observed that most transcripts of our input RNA pool were found in relatively few (<10%) condensates. 2. Only 0.1% of transcripts are found in almost all (<90%) of coacervates and 3. a substantial fraction of input transcripts (21%) were not detected in any sequenced condensate. This was likely due to low abundance of these transcripts in the input, although we cannot exclude a mechanism of exclusion due to currently unknown transcript features (Supplementary Fig. 3).

Next, we investigated which RNA features determine how frequently a transcript is found in coacervates. Generally, we found a strong relationship between input amount and the frequency of detection in coacervates (Fig. 3a). This indicates that the uptake of RNA is strongly dependent on the frequency the RNA is in the input. Interestingly, we found that there was a small subset of transcripts which did not follow this trend and were found in many or almost all of the coacervates, even though they were not very abundant in the input (Fig. 3a – red dots).

**Fig. 3.**
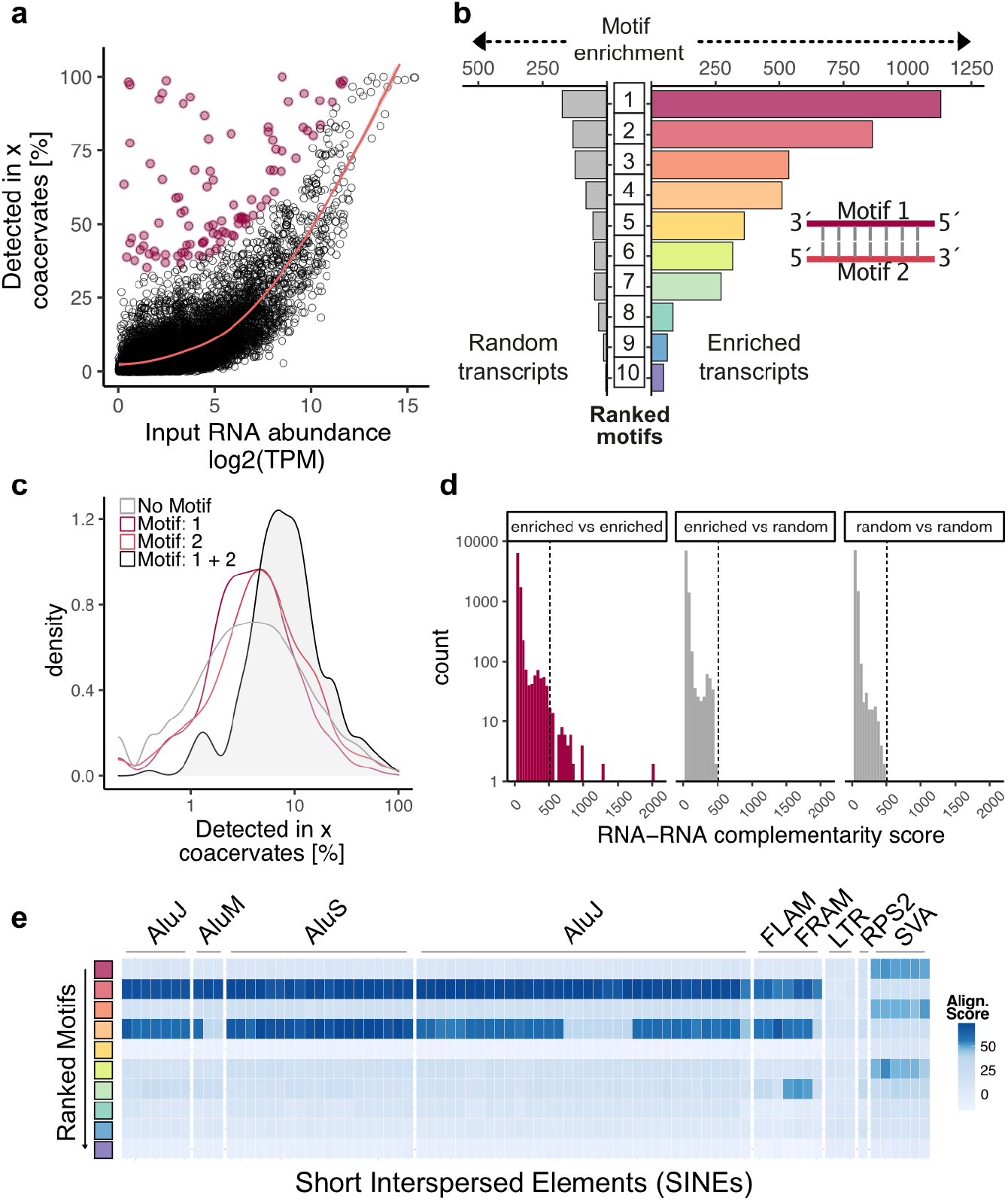
Properties of RNA found within coacervates. **a**, Correlation between input RNA amount and the frequency with which each transcript is detected in CM-Dex:PDDA coacervates. Transcripts that are enriched in coacervates (defined as residuals > 30 for generalized additive model) are labelled red. **b**, Analysis of sequence motifs that are detected within enriched transcripts (as defined in **a**) or randomly selected non-enriched transcripts. Among enriched transcripts, the two most abundant sequence motifs (Motifs 1 and 2) display sequence complementarity. **c**, Frequency of transcript detection in coacervates conditional on if the transcripts contain either motif 1, motif 2, both motifs or none. **d**, Analysis of sequence complementarity among different transcripts present in the pool of enriched or randomly selected transcripts. Sequence complementarity was determined using local-pairwise alignment (Smith-Waterman) scores. Dotted line indicates the maximum complementarity score that was detected outside the *enriched vs. enriched* comparisons (gray bars). **e**, Comparison of sequence similarity of enriched motifs to known genomic elements. Heatmap represents pairwise alignment (Smith-Waterman) of enriched motifs with sequences of short interspersed elements (SINEs). Color intensity represents alignment score.

We tested if other RNA features such as length or sequence might explain the efficient uptake of these transcripts. Our analysis showed that there was no correlation to the transcript length and its frequency in detection into the coacervates (Supplementary Fig. 4). However, sequence analysis of the RNA which were not highly abundant in the input but were frequently found within the coacervate droplets showed that there were sequence motifs of 11-50 bp which were enriched in the droplets compared to randomly selected non-enriched transcripts (Fig. 3b). Closer inspection of the sequence motifs revealed that the two most highly ranked motifs (Motif 1 and Motif 2) were in fact almost perfect reverse complements of each other (Fig. 3b and supplementary Fig. 5). To investigate the effect of Motif 1 and Motif 2 on transcript uptake by coacervates, we looked at the efficiency of uptake of an RNA transcript which contained both motifs. We found that transcripts which contained both motifs on the same transcript were detected more frequently within a coacervate compared to transcripts containing just Motif 1 or Motif 2 alone (Fig. 3c). The distances between these motifs on the transcripts were, however, too large to suggest hairpin structures (Supplementary Fig. 6a), potentially pointing towards more intricate secondary RNA structure. This is further supported by the fact that some motifs in enriched transcripts have very defined distances (median distances: 70, 71, 53, 84 for motifs 1, 4, 6 and 9 respectively) to each other when detected on the same transcript (Supplementary Fig. 7a,b) In contrast, all motifs found in randomly chosen transcripts displayed a broad distribution of distances to other motifs suggesting no obvious structural relationship between those motifs (Supplementary Fig. 7c,d). Since this result suggested that RNA-RNA interaction through sequence complementarity on the same transcript might be an important determinant of efficient RNA uptake into coacervates, we further investigated sequence complementarity across different transcripts. We found that the pool of enriched transcripts contains transcript pairs with very high sequence complementarity compared to enriched vs. random transcripts or random vs. random transcripts (Fig. 3d). In order to more directly test the impact of double-stranded RNA formation for uptake into coacervates we synthesized fluorescently-labelled oligonucleotides of Motif 1 and Motif 2 and quantified the uptake with flow cytometry. While quantifying a large number of coacervates (n = 10000), we observed that coacervates take up more double stranded RNA composed of Motif 1 and 2 compared to each motif alone or scrambled motifs (Supplementary Fig. 6b).

Next, we sequence matched the discovered sequence motifs to match any known genomic features. The motifs showed high similarity to genomic regions annotated as short interspersed elements (SINEs). SINEs belong to the family of transposable elements which have the potential to regulate transcription or generate new transcript isoforms^11^. In order to systematically test for sequence homology, pairwise alignment of each motif with SINE family members was undertaken (Fig. 3e). It was found that two motifs (motif 2 and 4) show strong sequence similarity to Alu elements which are primate specific transposable elements which are highly abundant in the human genome^12^. Three motifs (motif 1, 3 and 6) display similarly high homology to hominid-specific SINE-VNTR-Alu (SVA) retrotransposons which also has an Alu element as its main component^13^.

As the single cell sequencing methodology is applicable to both synthetic coacervate droplets and to coacervates which are formed from protein scaffolds we compared the RNA accumulation properties between different systems. We generated coacervates from CM-Dex with polylysine (CM-Dex:pLys, 6:1 molar ratio) to compare the results obtained so far to another synthetic coacervate system. Lysine residues are enriched in disordered regions of P-body condensate proteins and its polymer form has been shown to form condensates which support complex enzymatic reactions^14, 15^. Additionally, we sequenced RNA from well characterized Dhh1 and FUS-based phase separated droplets in order to compare RNA accumulation in synthetic coacervates versus protein-based condensates (Fig. 4a)^7, 16^.

**Fig. 4.**
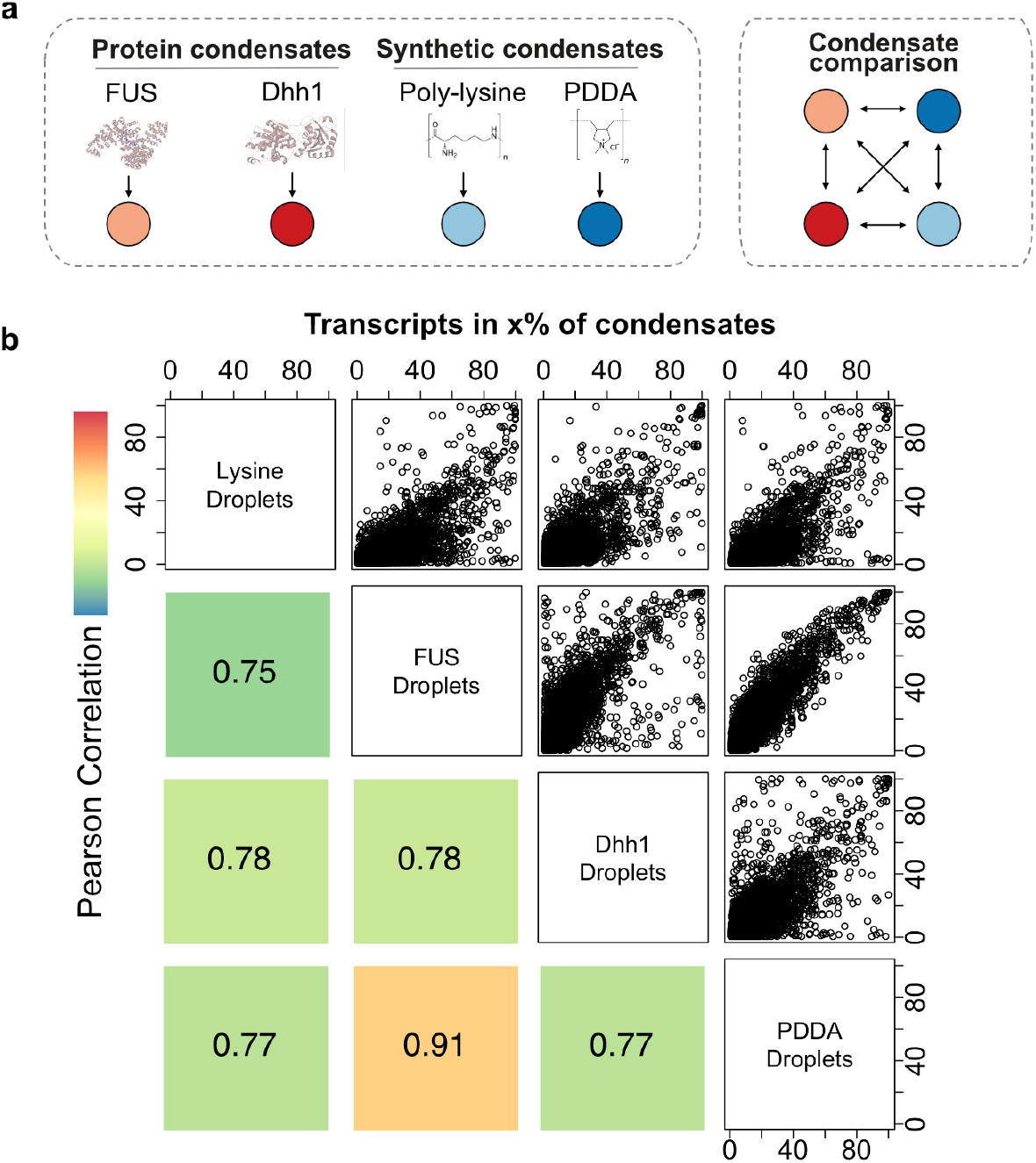
Comparison of RNA content across different coacervate and condensate types. **a**, Schematic representation of condensate types. Phase-separation of synthetic condensates (CM-Dex:PDDA, CM-Dex:pLys) was induced through addition of carboxlymethyldextran. **b**, Scatter plots and corresponding Pearson correlations comparing how frequently each transcript is detected in different condensate types. Color represents magnitude of correlation.

We first looked at how often any given transcript is detected in the different droplet systems and compared all results (Fig. 4b). We found a high correlation between all condensate types in particular for PDDA and FUS condensates (Fig. 4b). These results demonstrate that many RNAs that frequently localize in droplets will do so, irrespective of the host molecules of the droplets. However, there is a subpopulation of transcripts that are taken up more efficiently in a condensate-type specific way which was not a result of differences in the input (Fig. 4b and Supplementary Fig. 8).

For global cross comparison of all sequenced condensates we performed a dimensionality reduction analysis followed by Uniform Manifold Approximation and Projection (UMAP)^17^. This analysis evaluates how comparable all profiled condensates are to each other with respect to the RNA transcripts they contain. For this analysis, we focused on input independent, enriched transcripts for each condensate type (as defined in Fig. 3a) since we observed that there are many enriched transcripts that are specific to the chemical composition of the condensate types (Supplementary Fig. 10b). This also enables us to mitigate batch effects due to differences in the input RNA. We saw that FUS and PDDA condensates cluster closely together, whereas lysine condensates clustered with Dhh1 droplets indicating close RNA content similarity between these condensate types (Supplementary Fig. 9a). The Dhh1 condensates as well as the CM-Dex:pLys coacervates split into two clusters which are distinguished by condensate size indicating that small and large Dhh1 and CM-Dex:pLys droplets enrich for different transcripts (Supplementary Fig. 9b). We also performed motif enrichment analysis for all condensate types and found that the most enriched motif of the PDDA condensates was also highly enriched in all other condensate types (Supplementary Fig. 10a,c). Hence, this motif might confer advantages for transcripts to be taken up into condensates universally, irrespective of the molecular composition of the condensate.

## Discussion and conclusions

In summary, our data demonstrate for the first time that it is possible to explore the RNA content of single coacervate droplets. We dissected the molecular heterogeneity of a pool of coacervates allowing us to determine molecular differences between them. Thus far, differences between single coacervates could only be described on the phenotypic level by microscopy. Our ability to combine the sequencing data describing the RNA content with the FACS data describing the size and granularity of coacervates enabled us to link genotype and phenotype on the level of individual coacervates. Understanding the genotype-phenotype link is of primary importance towards the generation of artificial cells, the origin-of-life and for modern biology^19^.

A central question regarding the genotype of coacervates is, what types of RNAs it enriches for and which features the RNA molecules are characterized by. We found that RNAs with high sequence complementarity within or across RNA sequences are enriched in coacervates. This finding is reminiscent of the fact that stress granules, which form through liquid-liquid phase separation in cells, enrich ncRNAs which are complementary to mRNAs and likely form double stranded RNA^20^. Hence, increased charge density as a consequence of RNA double strand formation might be a prevalent feature of RNA content in biomolecular condensates which can be recapitulated in *in-vitro* reconstituted synthetic coacervate systems.

We further found that coacervates enrich for RNAs that contain sequence motifs that strongly resemble short interspersed elements (SINEs) and in particular Alu elements. Interestingly, Alu element-containing RNAs were previously shown to be enriched in the nucleolus, the largest condensate in the cell nucleus of eukaryotic cells^21, 22^. Our data therefore indicate that interactions of complementary Alu elements within transcripts could lead to formation of double stranded RNA. This interaction, rather than overall differences in global RNA structure (Supplementary Fig. 6c) likely represent a key RNA feature that leads to enriched RNA localization into coacervates.

When we compare the RNA content of protein-based condensate and synthetic polymer-based coacervates we found many similarities. Many transcripts that frequently enter one type of condensate also do so for others. Additionally, enriched transcripts for all condensate types are enriched for SINE sequence motifs, suggesting that these motifs confer an advantage to condensate localization irrespective of the molecular composition of the condensate type. These results demonstrate that synthetic coacervates represent a simple and cost-effective system for studying the physicochemical determinants of RNA localization during biomolecular condensation driven by proteins such as FUS or Dhh1.

Taken together, our data demonstrate that single cell RNA sequencing technology is not confined to the analysis of living cells but also applicable to RNA characterization of in-vitro phase separated coacervates^18^. It allows for highly multiplexed analysis of multiple condensate types and has the potential to uncover many aspects of the role of RNA in condensate formation with implications on several scientific disciplines from chemistry to cell biology.

## Acknowledgments

We would like to thank Malgorzata Santel, Anne Weigert and Theresa Schaffer for FACS sorting condensates. Further, we thank Barbara Schellbach and Antje Weihmann for performing Illumina sequencing. We also want to thank Hannes-Claudius Schulze for experimental help and Tobias Gerber, Sabina Kanton, Agnieska Brazovskaja, Stephan Bernhart, Jörg Fallmann and the Treutlein/ Camp labs for helpful discussions. Funding: This work was supported by core funds of the Max Planck Society and ETH Zurich, by ERC StG 758877 ORGANOMICS, the Volkswagen Foundation as well as by ERA-NET rare disease research implementing IRDiRC objectives - N° 643578 - Grant: REPETOMICS;

## Author contributions

D.W. performed the experiments. D.W. and B.V. performed data analysis. J.W. and A.H. provided recombinant FUS-GFP. M.H. and K.W. provided recombinant Dhh1-mCherry. D.W., G.C., T.-Y.D.T. and B.T. designed experiments and wrote the manuscript. Competing interests: Authors declare no competing interests.

## Supplementary information

**Fig. S1.**
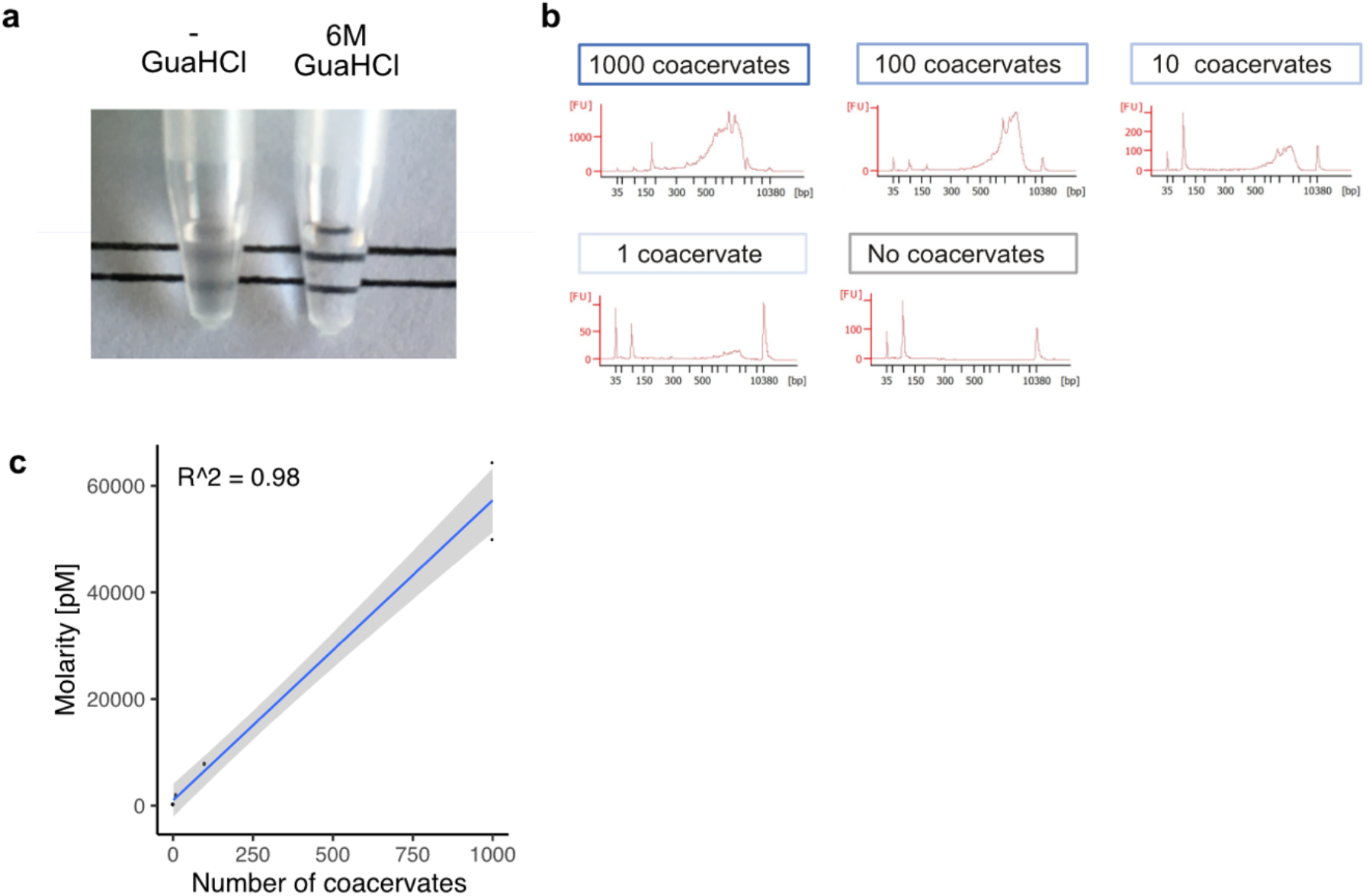
RNA extraction and library preparation from sorted coacervate droplets. a, Effect of 6M guanidine hydrochloride (GuaHCl) on the turbidity of PDDA:CM-Dex solution. **b**, Bioanalyzer traces for quantification and quality control of amplified cDNA prepared from multiple, single or no coacervates. **c**, Linear relationship between the number of coacervates (1000, 100, 10 1) sorted into a well and the resulting amplified cDNA library. **d**, Comparison of global coacervate similarity regarding selected parameters using principal component analysis (PCA). Numbers in brackets indicated how much the global variance across coacervates is explained by the respective components.

**Fig. S2.**
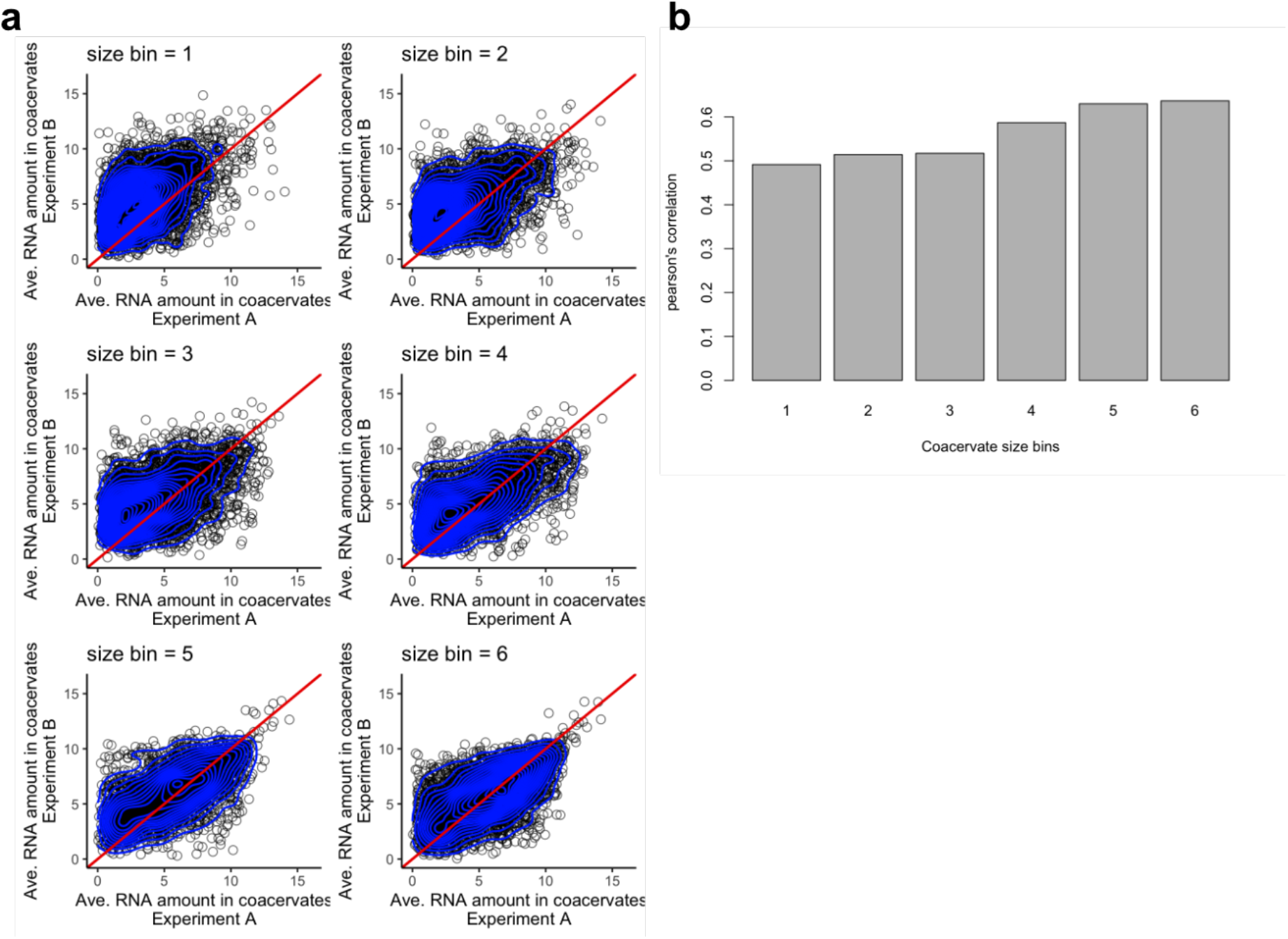
Effect of droplet size on consistency of the number of transcripts found in coacervates. **a**, Experiment-to-experiment variation of the average abundance of each RNA transcript across all CM-Dex:PDDA coacervates in which it was detected as in Fig. 2b. Each plot represents data for a subset of coacervates (size bin) of a given size range. Size bin 1 refers to the smallest and 6 to the largest coacervates. All size bins are of equal size regarding the number of coacervates they contain. **b**, Comparison of pearson’s correlations for all size bins of Fig. S2a.

**Fig. S3.**
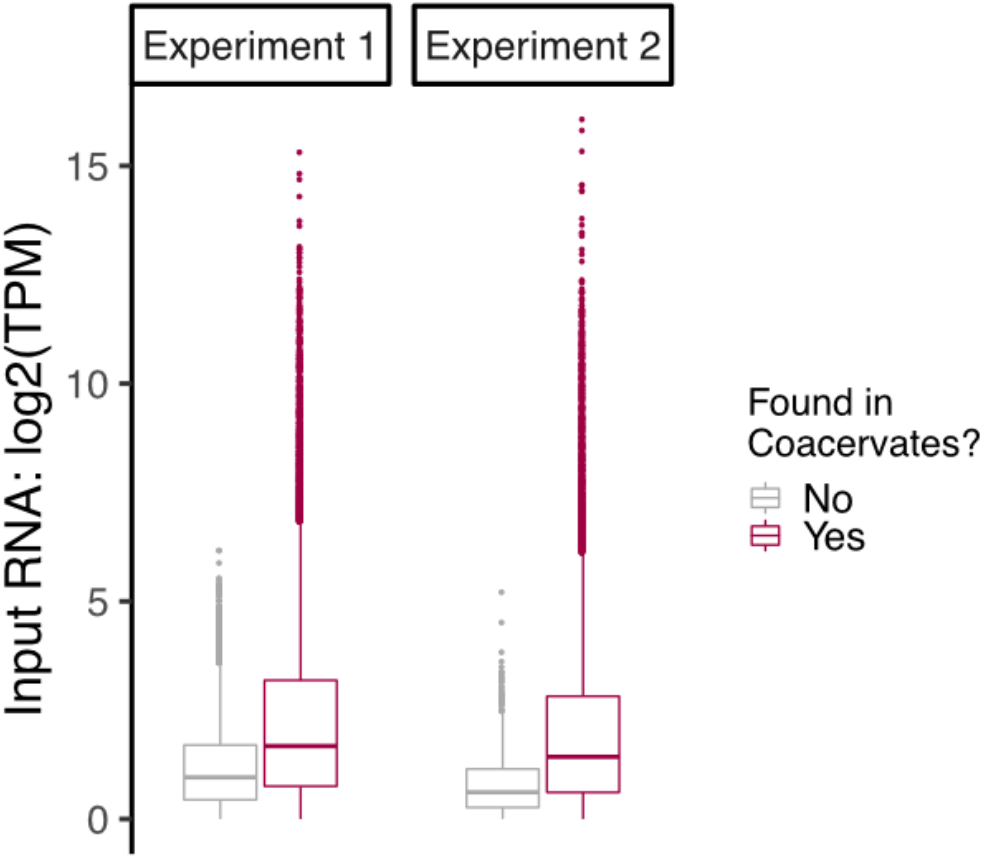
Transcripts not detected in coacervates were not abundant in the input RNA pool. Relationship between transcript abundance in the input of each experiment and whether it was detected in at least one CM-Dex:PDDA coacervate in the respective dataset. Transcript abundance in the input was measured as transcripts per kilobase million (TPM).

**Fig. S4.**
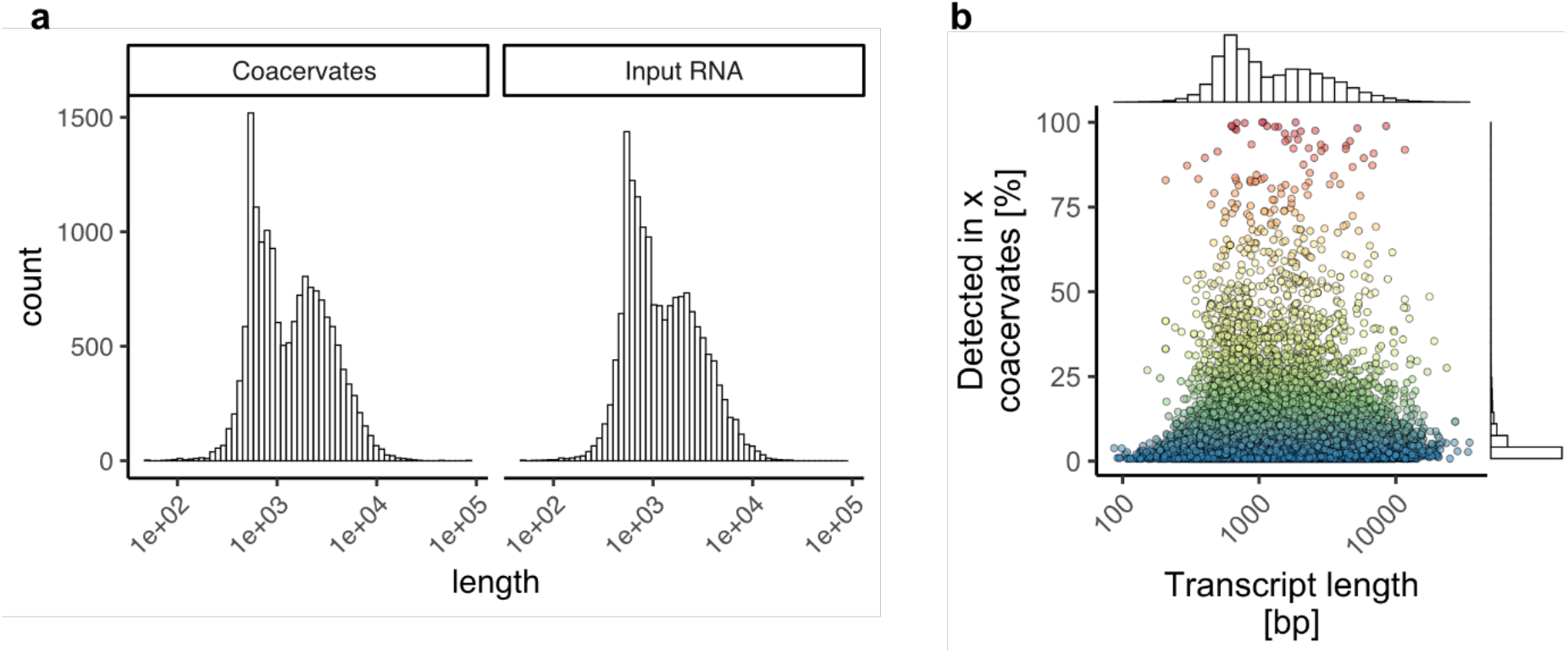
The effect of transcript length on RNA partitioning into coacervates. **a**, Comparison of transcript lengths of all detected RNA transcripts in CM-Dex:PDDA coacervates and the input. **b**, Analysis of frequency of transcript detection in coacervates as a function of transcript length.

**Fig. S5.**
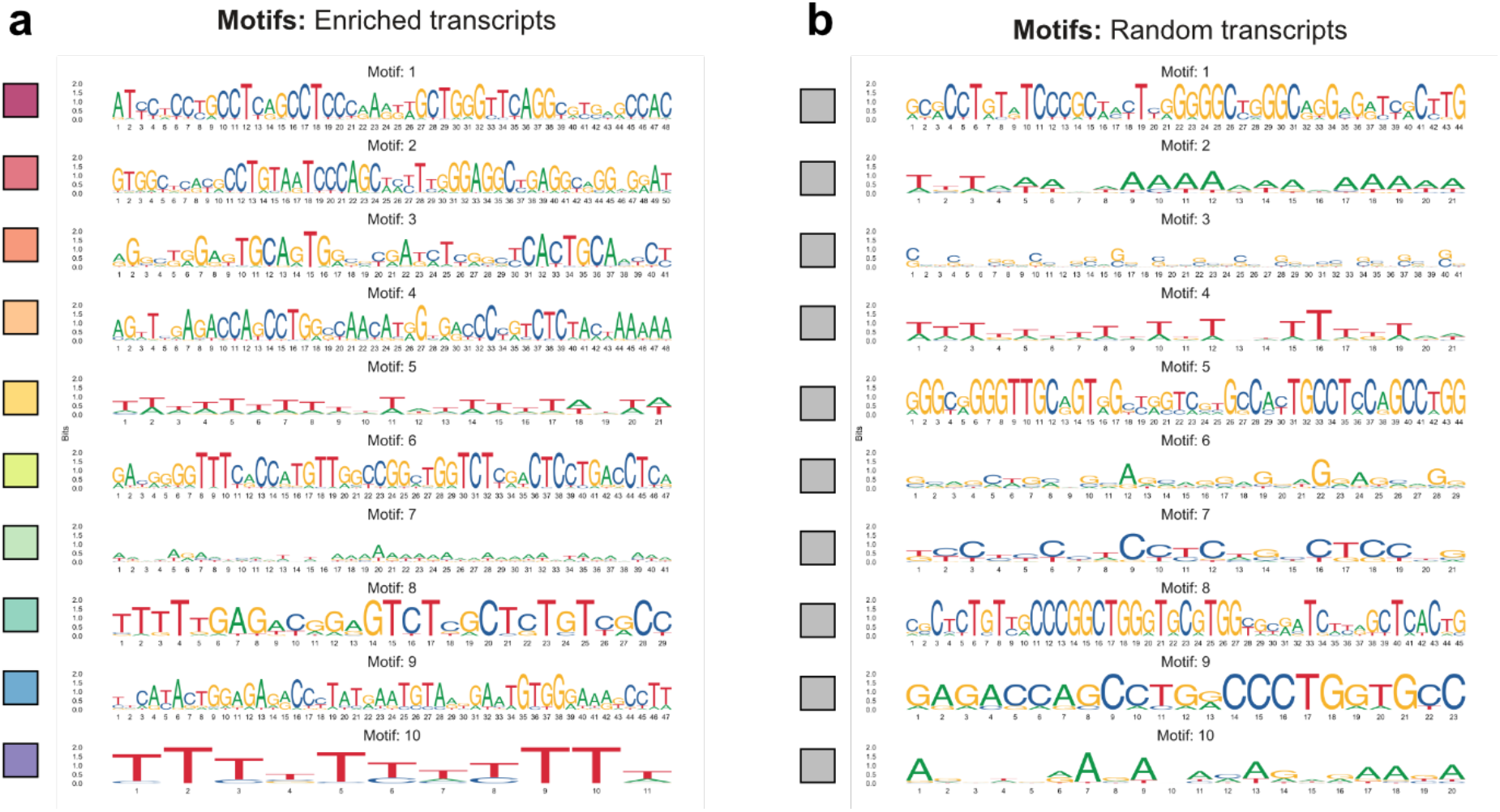
Top motifs in enriched and randomly selected non-enriched transcripts. **a**, Sequences of top 10 motifs (ranked descending from top to bottom) in enriched transcripts (as defined in Fig. 3a) and the same number of randomly selected non-enriched transcripts. **b**, Motif enrichment values (MEME E-value) displayed in Fig. 3b. Colors of squares represent transcript pool (enriched vs. random) and motif enrichment as in Fig. 3b

**Fig. S6.**
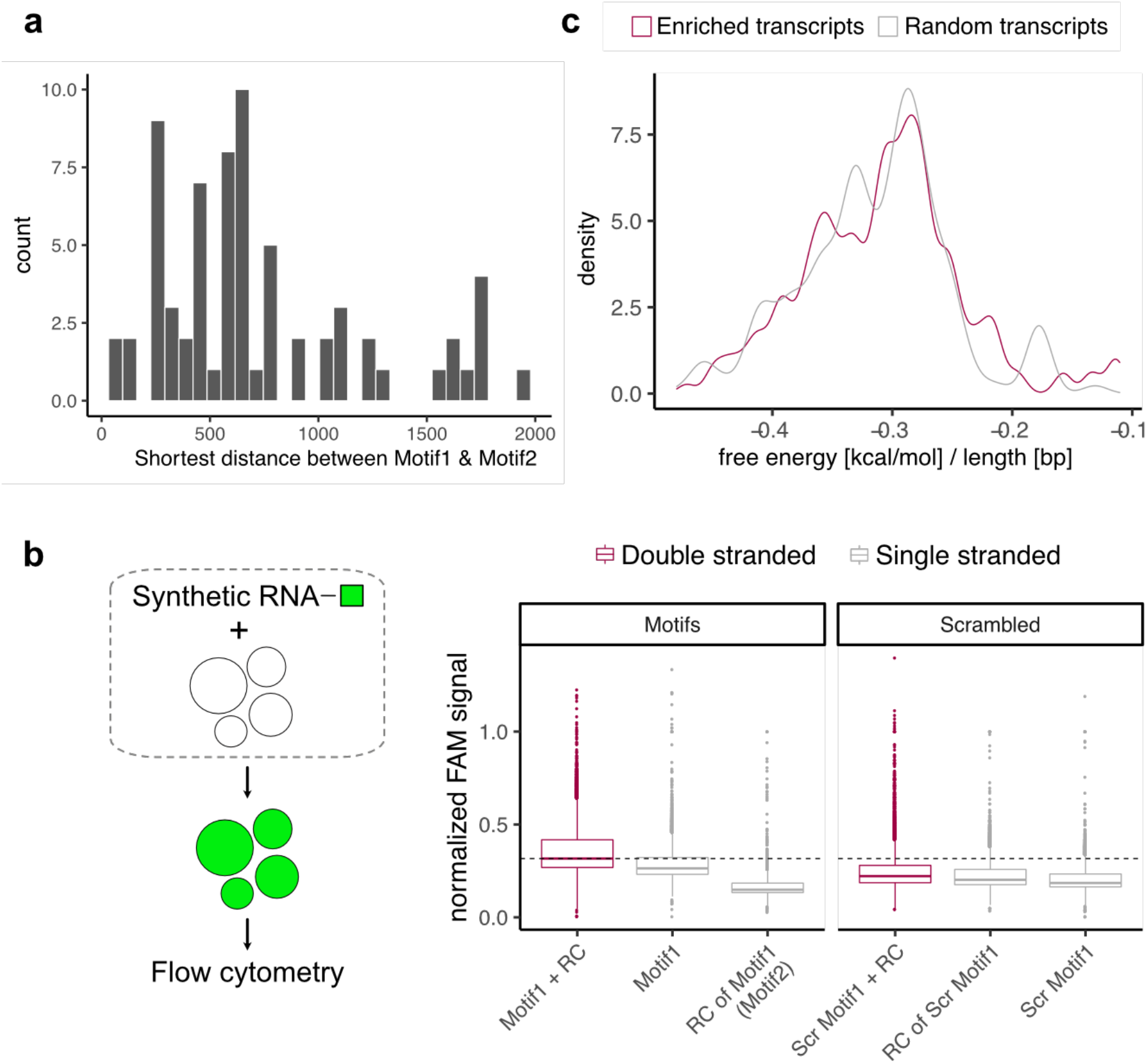
Relationship between the two most abundant motifs and RNA folding analysis. **a**, Distribution of distances between the two most enriched motifs (Motif1 and Motif2) found among enriched transcripts (see Fig 3a,b). For potential hairpin formation, only the shortest distances between the motif were considered for each transcript. **b**, Quantification of CM-Dex:PDDA coacervate uptake of different chemically synthesized sequences. Coacervate uptake of FAM-labelled oligonucleotides were analyzed by flow cytometry. *Motif 1* (most enriched motif - see Fig. 3b and Supplementary Fig. 2), *Motif 2* (its reverse complement (RC)), *Scrambled Motif 1* (scrambled sequence of Motif 1) and the reverse complement (RC) of *Scrambled Motif 1* were analyzed. *Double stranded* refers to pre-mix of Motif 1 or Scrambled Motif 1 with their respective reverse complement. **c**, Comparison of the minimum free energies (normalized for transcript length) of enriched and randomly selected transcripts

**Fig. S7.**
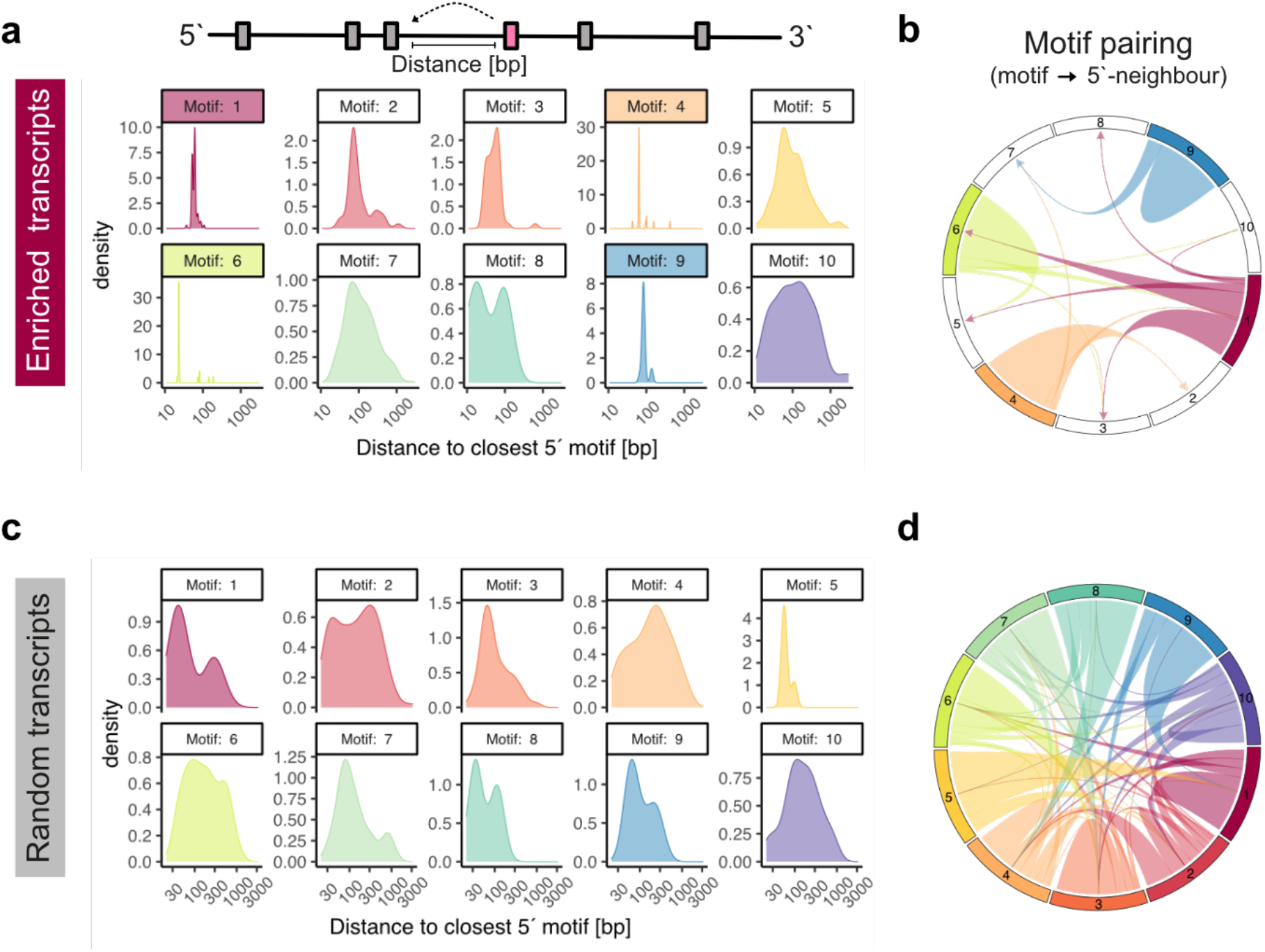
Sequence motif distances on transcripts. **a**, Distribution of distances between each motif (detected in enriched transcripts) and its closest 5`neighbour on a given transcript. Colored facets highlight motifs with narrow distributions. **b**, Circos plot depicting how many times each highlighted motif (see Fig. S6A) pairs with the other motifs as their respective 5` neighbor. **c**, Distribution of distances between each motif (detected in randomly selected transcripts) and its closest 5`neighbor. **d**, Circos plot depicting how many times each motif pairs with the other motifs as their respective 5` neighbor.

**Fig. S8.**
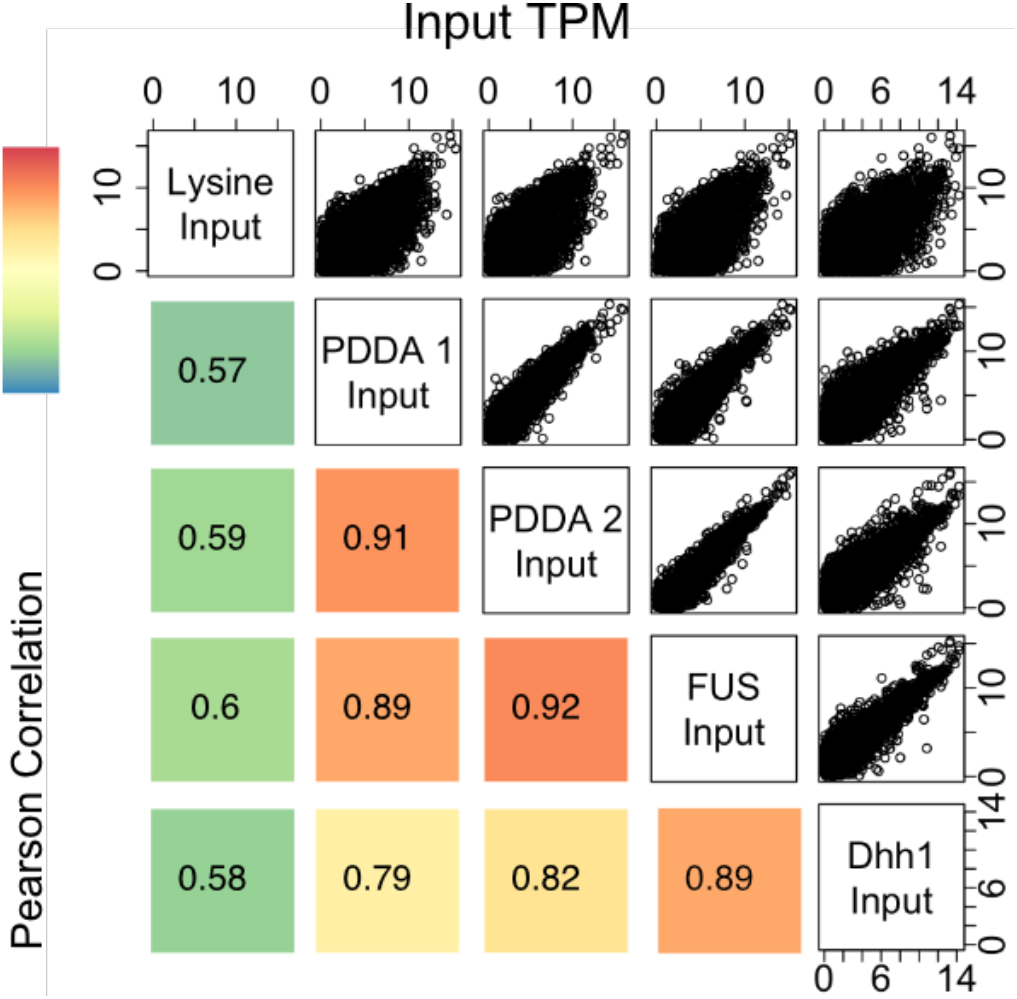
Experiment-to-experiment variation of input RNA abundances. Scatter plots and corresponding Pearson correlations comparing the abundances of all input transcripts (log_2_(TPM)) across different experiments and condensate types. Colors represent magnitude of correlation.

**Fig. S9.**
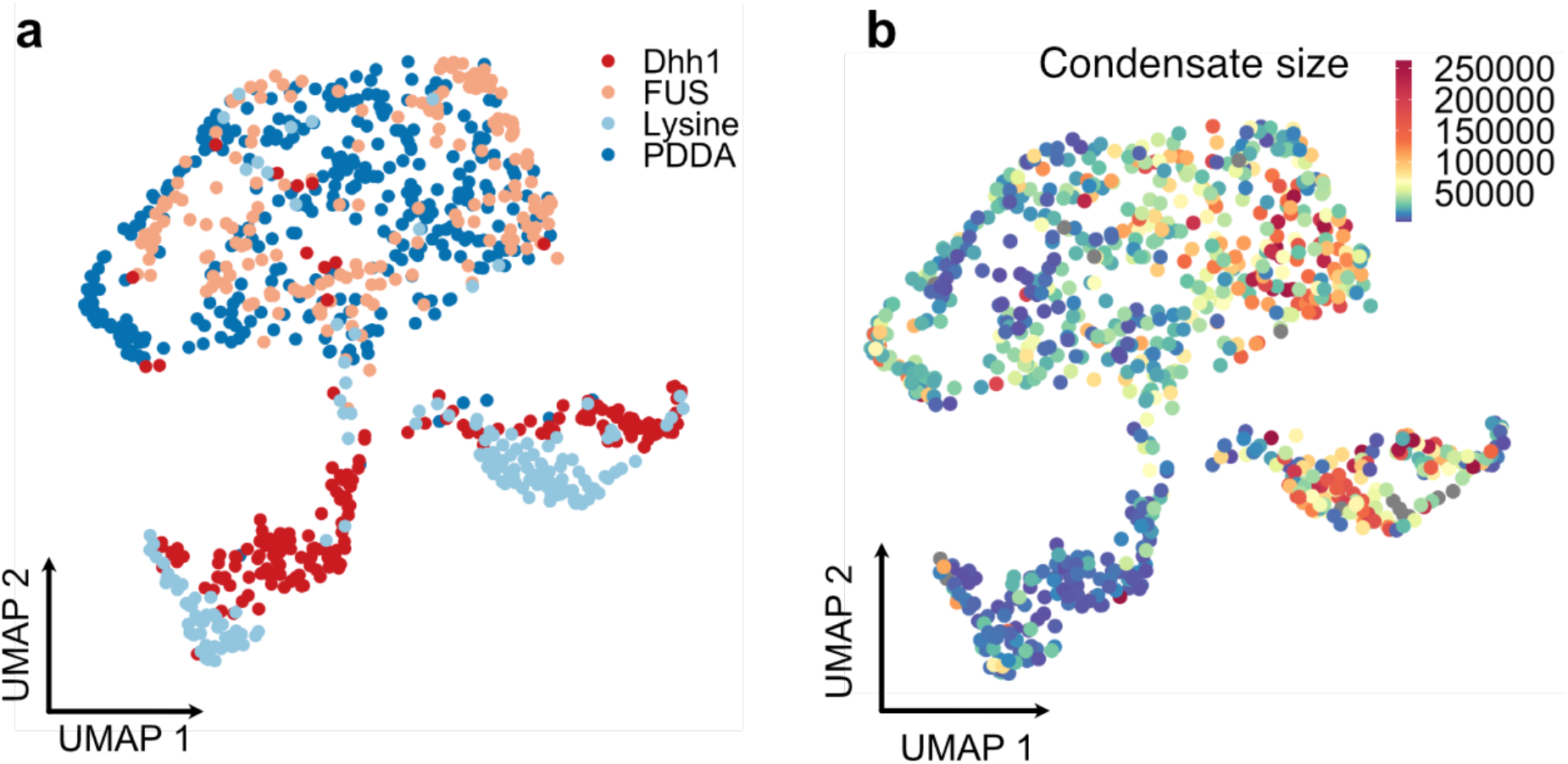
Global comparison of RNA content across all condensate types. **a**, Uniform Manifold Approximation and Projection (UMAP) analysis reduces the dimensionality of the data in order to visualize condensate similarities and differences across thousands of genes. Each dot represents a condensate. Colors represent different condensate types. **b**, Same UMAP as in a with color code representing the size of the condensate as measured by FACS. Legend values correspond to forward-scatter (FSC) values obtained from FACS analysis.

**Fig. S10.**
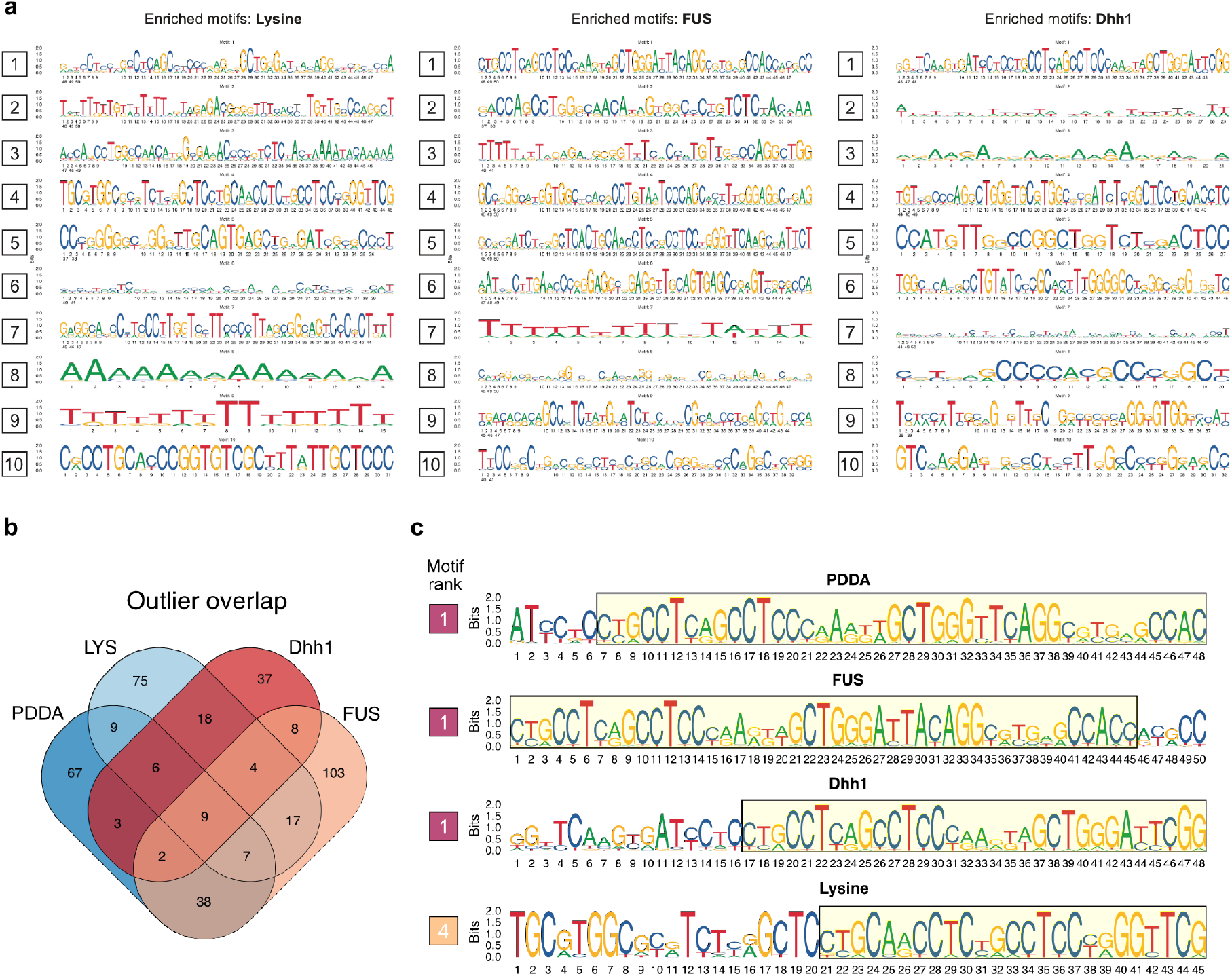
Sequence motif analysis of enriched transcripts across condensate types. **a**, Sequences of top 10 motifs (ranked descending from top to bottom) detected in enriched transcripts of Lysine-CM-Dextran, FUS or Dhh1 droplets. **b**, Quantification of overlap between transcript that are enriched (residuals > 30 - see Fig 3a) in different condensate types. **c**, Comparison of motif similarity between the top motif of the PDDA condensates and sequence similar motifs found in enriched transcripts of each condensate type.

**Table S1.**
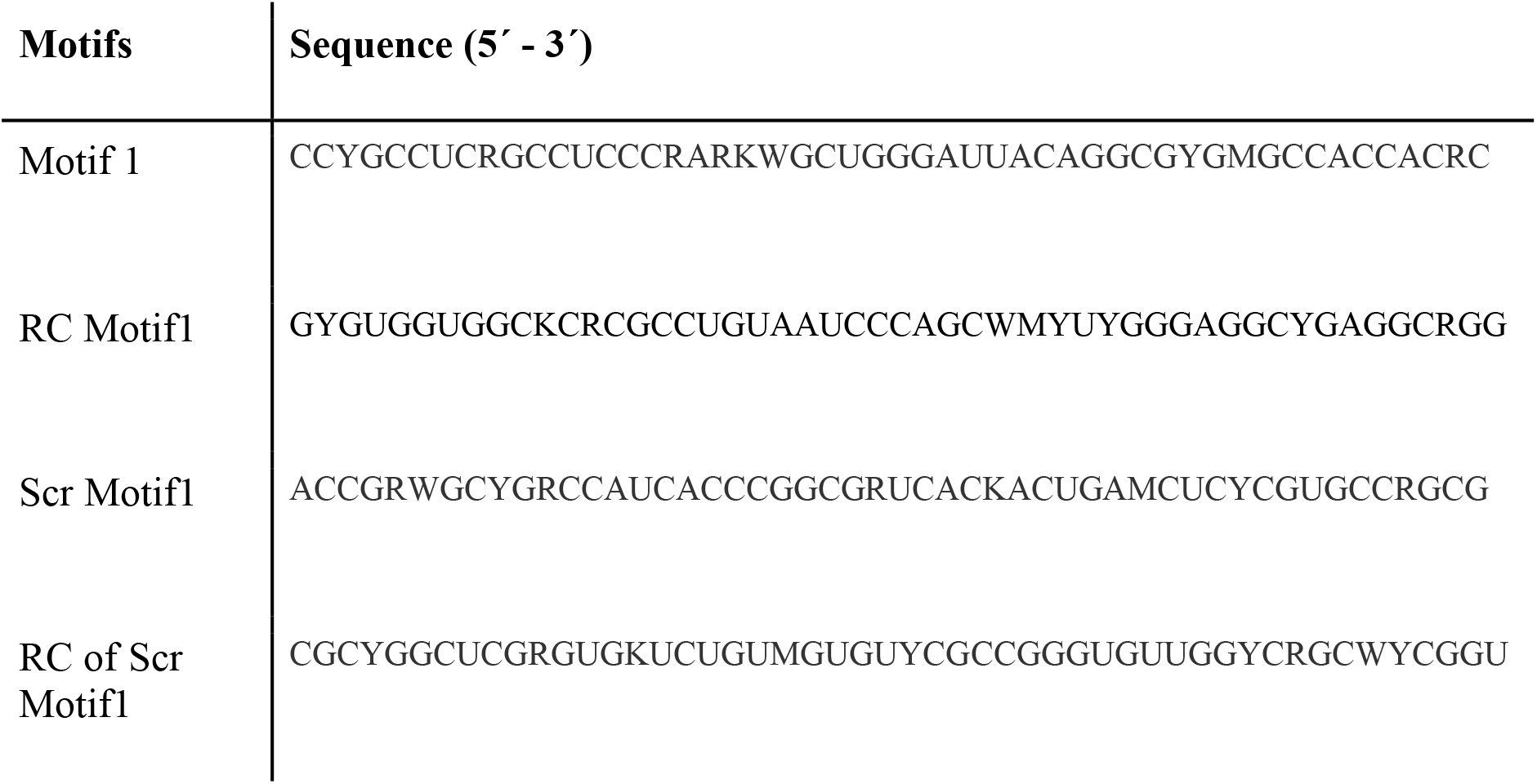
Chemically synthesized motif sequences.

**Table S2.**
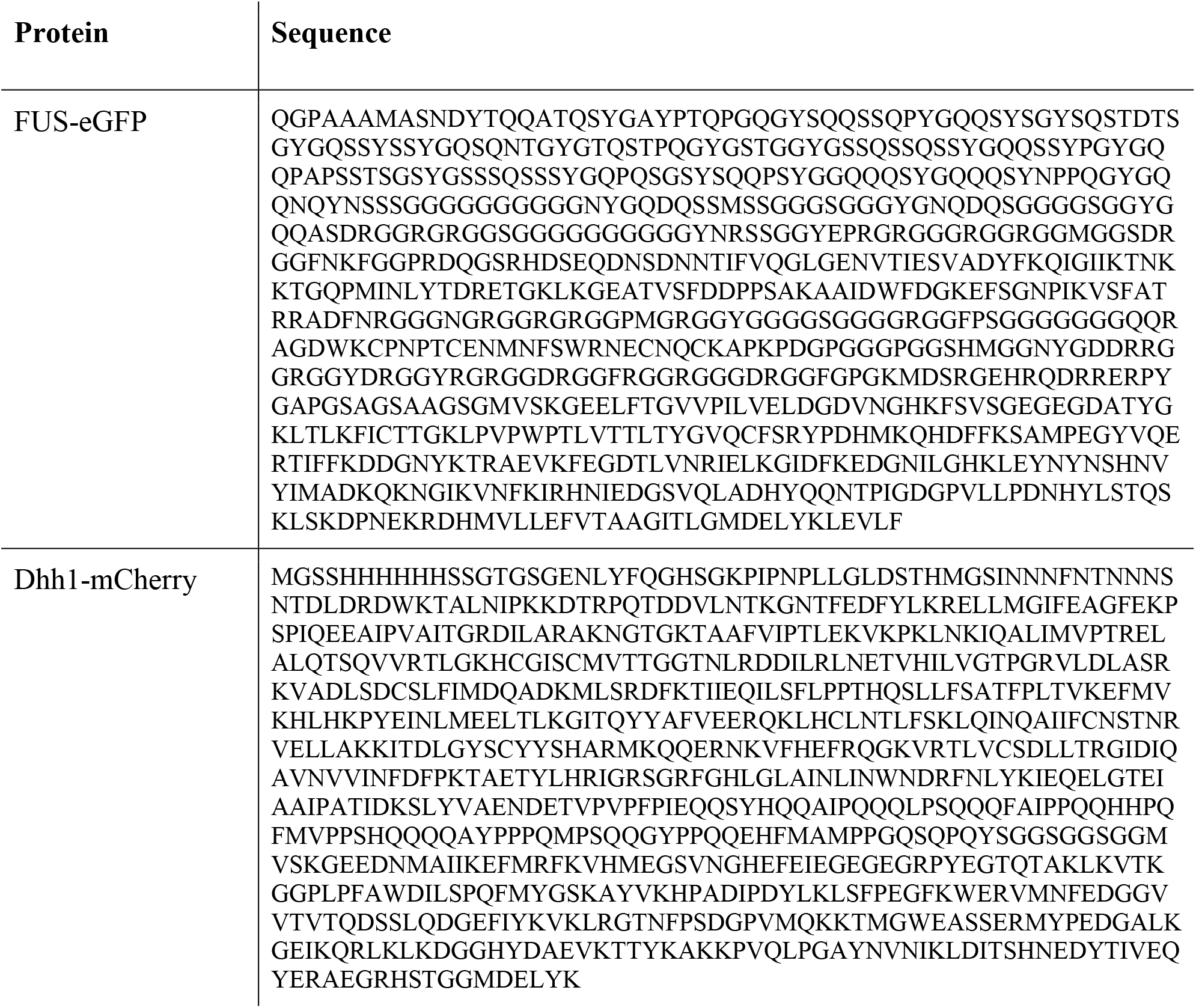
Amino acid sequence of recombinant proteins.

## Materials and Methods

### Condensate generation

Synthetic polymer-based coacervates were prepared as previously described (*32*). Specifically, Poly(Diallyl Dimethyl Ammonium Chloride) (PDDA, 8.5 kDa, monomer: 161.8 g·mol^−1^) or poly-L-lysine (4–15 kDa, monomer: 161.67 g mol^−1^) were mixed with CM-Dex sodium salt (10–20 kDa, monomer: 162.14 g mol^−1^) at a molar ratio of 6:1 in Tris-MgCl_2_ buffer (10mM Tris-HCl pH 8.0 and 4mM MgCl_2_). Total RNA was isolated from iPSC cells (409B2) using the RNeasy Mini Kit (Qiagen). Coacervates were generated by adding CM-Dex, RNA and PDDA in the respective order to the Tris-MgCl_2_ buffer to achieve a final RNA concentration of 50 ng/μl. Coacervates were then incubated for 1h at room temperature while rotating before FACS sorting. FUS-GFP and Dhh1-mCherry proteins were cloned, purified and respective protein-based condensates were prepared as previously described (*4, 30*). Recombinant FUS-GFP in 25 mM Tris-HCl pH 7.4, 150 mM KCl, 2.5% Glycerol and 0.5 mM DTT was used at a final protein concentration of 1 mg ml^−1^. Dhh1-mCherry in 200 mM NaCl, 25 mM Tris (pH 7.4) and 10% glycerol was used at a final protein concentration of 150 μM. Dhh1 droplets were generated by adding ATP (final conc. 10 mM), creatin kinase based ATP recombination system CKM (40 mM ATP, 40 mM MgCl2, 200 mM creatine phosphate, 70 U/mL Creatine Kinase), BSA (final conc. 1mg ml^−1^) and Hepes buffer (final conc. 50 mM) to the recombinant Dhh1-mCherry protein in low salt buffer (50 mM KCl, 30 mM HEPES-KOH pH 7.4, 2 mM MgCl_2_).

### Single coacervate index sorting

RNA containing coacervates (initial volume min. 250 μl) were sorted with a BD FACSAria Fusion flow cytometer using a 150 μl nozzle. Single-coacervates were index-sorted in precooled skirted twin.tec 96-well LoBind Plates (Eppendorf) containing 4 μl of 6 M Guanidine HCl (GuaHCl, Sigma) as lysis buffer. For each plate, one well was sorted with 1000 coacervates and one well was left empty as positive and negative controls respectively. Directly after sorting the plates were briefly spun down (max speed) to collect all FACS-derived droplets in the lysis buffer. The plates were then immediately put on dry ice until all other plates were sorted. Plates were kept at −80 °C until cDNA was prepared.

### Bulk coacervate FACS analysis

Coacervates with and without RNA were analysed on a BD FACSAria Fusion (150 mm nozzle) and data was processed in R using the *flowcore* package. For quantification of RNA incorporation into coacervates, size-matched FAM-labelled RNAs were synthesized (IDT) and incorporated into PDDA-CM-Dex coacervates as described above. Sequences of chemically synthesized oligos can be found in Table S1.

### Single coacervate library preparation

Before library preparation the plates were spun down to collect all liquid at the bottom of the wells. SPRI beads (Agencourt RNAclean XP, Beckman Coulter) were equilibrated to room temperature and 2.2x SPRI beads were added to each well. Upon incubation for 5 min at room temperature and beads were washed twice using 80% EtOH as described in the SPRI bead manufacturer’s protocol. EtOH traces were completely removed and beads were dried for 2-3 minutes (Note: Beads dry out fast after exposure to GuaHCl. Overdrying of beads will lead to significantly lower yields). RNA was eluted by resuspending beads in 3μl of dNTP/oligodT mix, then beads were magnetically separated from RNA/dNTP/oligodT mix and transferred to a new 96-well plate. Next, the SMART-seq2 protocol described in Picelli et al., 2014^10^ was followed from step 9 onwards with the following modifications: a) the template switching oligo was biotinylated on the 5-end b) PCR preamplification was performed for 23 cycles. Size distribution of cDNA obtained from single coacervates was checked for randomly chosen samples to verify success of cDNA preparation. Next, tagmented libraries were prepared and sequenced (100bp paired-end reads) on an Illumina HiSeq 2500 as described^10^.

### Data processing, quality control and analysis

Raw sequencing data was processed using custom scripts and aligned to reference human transcriptome (hg38 sourced from Ensembl) using Kallisto (v0.44.0) with standard parameters including - pseudobam flag to obtain read coverage across each transcript. Transcripts TPM values < 1 were filtered out. For datasets with low average low average pseudoalignment (<40%), transcripts with less than 20% read coverage were excluded. Furthermore, since the coacervate size correlates with the number of transcripts detected as a consequence of coacervate size-dependent RNA concentrations we filtered out coacervates with < 5% pseudoalignment for sizes FSC > 2e4. Enriched transcripts (Fig. 3a) were defined as transcripts whose residuals value was > 30 when the data was fitted to a generalized additive model.

### Motif enrichment analysis

De novo motif discovery was determined using MEME (v5.0.5) with the following parameters: *-dna -time 18000 -mod anr -nmotifs 10 -minw 6 -maxw 50 -objfun classic -markov_order 2*. Sequences obtained from the reference human transcriptome (hg38 sourced from Ensembl) were chosen as input for MEME analysis. The background was calculated using the sequences of all input transcripts. Enrichment of discovered motifs for each transcript was calculated using MAST (v5.0.5) with *-nostatus -minseqs 21978 -remcorr -sep -ev 0.05 -c 1* parameters. MEME and MAST outputs were parsed for analysis in R using custom python scripts. Distances between every detected motif and its closest 5’-neighbour were calculated for each enriched transcript (from motif-start to motif-start – Fig. S6). The same analysis was done for motifs detected in randomly non-enriched transcripts (Fig. S6). Data was plotted using the *ggplot2* and *circlize* packages.

### RNA folding analysis

Analysis of minimum free energy for each enriched transcript and the same number of randomly selected non-enriched transcripts was performed using RNAfold (v2.4.12) with standard settings. RNAfold output was parsed for analysis in R using a custom python script.

### Comparison of sequence complementarity

The presence of Motif 1 and its reverse complement Motif 2 on the same transcript (*cis* complementarity – Fig. 3d) was determined using MAST with E-values < 0.05 as a cutoff. The complementarity of sequences across different transcripts (*trans* complementarity – Fig. 3d) was obtained by determining the pairwise local alignment using the Smith-Waterman algorithm. Briefly, two pools of transcripts were used for this analysis: enriched transcripts (residuals > 30 – see Fig. 3a) and the same number of randomly chosen non-enriched transcripts (residuals < 30) with a similar transcript length distribution. Local pairwise alignments for each transcript pair of the respective pools were calculated using the *Biostrings* package in R with the following parameters: *nucleotideSubstitutionMatrix(match = 2, mismatch = −1, baseOnly = TRUE), pairwiseAlignment(gapOpening = −30, gapExtension = −0.05, scoreOnly = TRUE, type=“local”)*. For comparison of enriched sequence motifs with SINEs (Fig. 3e) we obtained SINE reference sequences from RepBase (latest update: 08-24-2020). For each of the 10 consensus motifs, the 5 most significant motif hits found among the enriched transcripts were compared to each SINE sequence by pairwise alignment using the *Biostrings* R package. Then the pairwise alignment score was averaged over the 5 most significant motifs for each consensus motif providing the alignment score displayed in the plot. Alignment parameters: *nucleotideSubstitutionMatrix(match = 2, mismatch = −1, baseOnly = TRUE); pairwiseAlignment(gapOpening = −10, gapExtension = 0, scoreOnly = TRUE)*.

### Dimensionality reduction analysis

Principal component analysis (PCA) in Fig. S1d was performed using the *factoextra* R package. We used scaled coacervate morphology and transcript characteristics obtained from sequencing and FACS data as PCA features instead of relative transcript abundance in order to describe coacervate heterogeneity. Analysis of transcript-based coacervate heterogeneity (in Fig. S8) was conducted using the *Seurat* R package (v3.1.5). For this analysis, we used transcripts as input which were enriched for each coacervate type as defined by the analysis in Fig. 3a (residuals > 30 when fitted to a generalized additive model). These transcripts were normalized to the TPM values of the transcripts detected in the sequenced input RNA pool and subsequently scaled within each experiment. For clustering and UMAP analysis the first 5 principal components were used.

## References

1. de Jong, B. & Kruypz, H.R. Coacervation (partial miscibility in colloid systems). Proc. K. Ned. Akad. Wet. 32, 849–856 (1929).

2. Veis, A. A review of the early development of the thermodynamics of the complex coacervation phase separation. Adv Colloid Interfac 167, 2–11 (2011).

3. Nakashima, K.K., Vibhute, M.A. & Spruijt, E. Biomolecular Chemistry in Liquid Phase Separated Compartments. Front Mol Biosci 6(2019).

4. Oparin, A.I. The origin of life on the earth. Oliver & Boyd, Edinburgh & London 3rd Ed (1957).

5. Blocher, W.C. & Perry, S.L. Complex coacervate-based materials for biomedicine. Wires Nanomed Nanobi 9(2017).

6. Pak, C.W. et al. Sequence Determinants of Intracellular Phase Separation by Complex Coacervation of a Disordered Protein. Molecular Cell 63, 72–85 (2016).

7. Wang, J. et al. A Molecular Grammar Governing the Driving Forces for Phase Separation of Prion-like RNA Binding Proteins. Cell 174, 688-+ (2018).

8. Alberti, S. et al. A User’s Guide for Phase Separation Assays with Purified Proteins. Journal of Molecular Biology 430, 4806–4820 (2018).

9. Beneyton, T., Love, C., Girault, M., Tang, T.Y.D. & Baret, P.J.C. High-Throughput Synthesis and Screening of Functional Coacervates Using Microfluidics. ChemSystemsChem (2020).

10. Picelli, S. et al. Full-length RNA-seq from single cells using Smart-seq2. Nat Protoc 9, 171–181 (2014).

11. Elbarbary, R.A., Lucas, B.A. & Maquat, L.E. Retrotransposons as regulators of gene expression. Science 351(2016).

12. Deininger, P. Alu elements: know the SINEs. Genome Biology 12(2011).

13. Wang, H. et al. SVA elements: A hominid-specific retroposon family. Journal of Molecular Biology 354, 994–1007 (2005).

14. Tang, T.Y.D., van Swaay, D., deMello, A., Anderson, J.L.R. & Mann, S. In vitro gene expression within membrane-free coacervate protocells. Chem Commun 51, 11429–11432 (2015).

15. Ukmar-Godec, T. et al. Lysine/RNA-interactions drive and regulate biomolecular condensation. Nature Communications 10(2019).

16. Hondele, M. et al. DEAD-box ATPases are global regulators of phase-separated organelles. Nature 573, 144-+ (2019).

17. McInnes, L. & Healy, J. UMAP: uniform manifold approximation and projection for dimension reduction. Preprint at https://arxiv.org/abs/1802.03426 (2018).

18. Wollny, D. Opportunities for Single-Cell Sequencing in Synthetic Biology. ChemSystemsChem (2020).

19. Nourian, Z. & Danelon, C. Linking Genotype and Phenotype in Protein Synthesizing Liposomes with External Supply of Resources. Acs Synth Biol 2, 186–193 (2013).

20. Van Treeck, B. et al. RNA self-assembly contributes to stress granule formation and defining the stress granule transcriptome. P Natl Acad Sci USA 115, 2734–2739 (2018).

21. Caudron-Herger, M. et al. Alu element-containing RNAs maintain nucleolar structure and function. Embo Journal 34, 2758–2774 (2015).

22. Caragine, C.M., Haley, S.C. & Zidovska, A. Nucleolar dynamics and interactions with nucleoplasm in living cells. Elife 8(2019).

